# Isolation and characterization of SARS-CoV-2 from the first US COVID-19 patient

**DOI:** 10.1101/2020.03.02.972935

**Authors:** Jennifer Harcourt, Azaibi Tamin, Xiaoyan Lu, Shifaq Kamili, Senthil Kumar. Sakthivel, Janna Murray, Krista Queen, Ying Tao, Clinton R. Paden, Jing Zhang, Yan Li, Anna Uehara, Haibin Wang, Cynthia Goldsmith, Hannah A. Bullock, Lijuan Wang, Brett Whitaker, Brian Lynch, Rashi Gautam, Craig Schindewolf, Kumari G. Lokugamage, Dionna Scharton, Jessica A. Plante, Divya Mirchandani, Steven G. Widen, Krishna Narayanan, Shinji Makino, Thomas G. Ksiazek, Kenneth S. Plante, Scott C. Weaver, Stephen Lindstrom, Suxiang Tong, Vineet D. Menachery, Natalie J. Thornburg

## Abstract

The etiologic agent of the outbreak of pneumonia in Wuhan China was identified as severe acute respiratory syndrome associated coronavirus 2 (SARS-CoV-2) in January, 2020. The first US patient was diagnosed by the State of Washington and the US Centers for Disease Control and Prevention on January 20, 2020. We isolated virus from nasopharyngeal and oropharyngeal specimens, and characterized the viral sequence, replication properties, and cell culture tropism. We found that the virus replicates to high titer in Vero-CCL81 cells and Vero E6 cells in the absence of trypsin. We also deposited the virus into two virus repositories, making it broadly available to the public health and research communities. We hope that open access to this important reagent will expedite development of medical countermeasures.

**Article Summary:** Scientists have isolated virus from the first US COVID-19 patient. The isolation and reagents described here will serve as the US reference strain used in research, drug discovery and vaccine testing.

## BACKGROUND

A novel coronavirus, severe acute respiratory syndrome coronavirus 2 (SARS-CoV-2), has been identified as the source of a pneumonia outbreak in Wuhan China in late 2019 (1, 2). The virus was found to be a member of the beta coronavirus family, in the same species as SARS-CoV and SARS-related bat CoVs (3, 4). Patterns of spread indicate that SARS-CoV-2 can be transmitted person-to-person, and may be more transmissible than SARS-CoV (5–7). The spike protein of coronaviruses mediates virus binding and cell entry. Initial characterization of SARS-CoV-2 spike indicate that it binds the same receptor as SARS-CoV, ACE2, which is expressed in both upper and lower human respiratory tracts (8). The unprecedented rapidity of spread of this outbreak represents a critical need for reference reagents. The public health community requires viral lysates to serve as diagnostic references, and the research community needs virus isolates to test anti-viral compounds, develop new vaccines, and perform basic research. In this manuscript, we describe isolation of virus from the first US COVID-19 patient and described its genomic sequence and replication characteristics. We have made the virus isolate available to the public health community by depositing into two virus reagent repositories.

## RESULTS and DISCUSSION

A patient was identified with confirmed COVID-19 in Washington State on January 22, 2020 with cycle threshold (C_t_s) of 18-20 (nasopharyngeal(NP)) and 21-22 (oropharyngeal (OP)) (1). The positive clinical specimens were aliquoted and refrozen inoculation into cell culture on January 22, 2020. We first observed cytopathic effect (CPE) 2 days post inoculation and harvested viral lysate on day 3 post inoculation (Figure 1B and 1C). Fifty μl of P1 viral lysates were used for nucleic acid extraction to confirm the presence of SARS-CoV-2 using the CDC molecular diagnostic assay (1). The Cts of three different nucleic acid extractions ranged from 16.0-17.1 for N1, 15.9-17.1 for N2 and 16.2-17.3 for N3, confirming isolation of SARS-CoV-2. A C_t_ of less than 40 is considered positive. The extracts were also tested for the presence of 33 additional different respiratory pathogens with the fast track 33 assay. No other pathogens were detected. Identity was additionally supported by thin section electron microscopy (Figure 1D). We observed a morphology and morphogenesis characteristic of coronaviruses.

**Figure 1.**
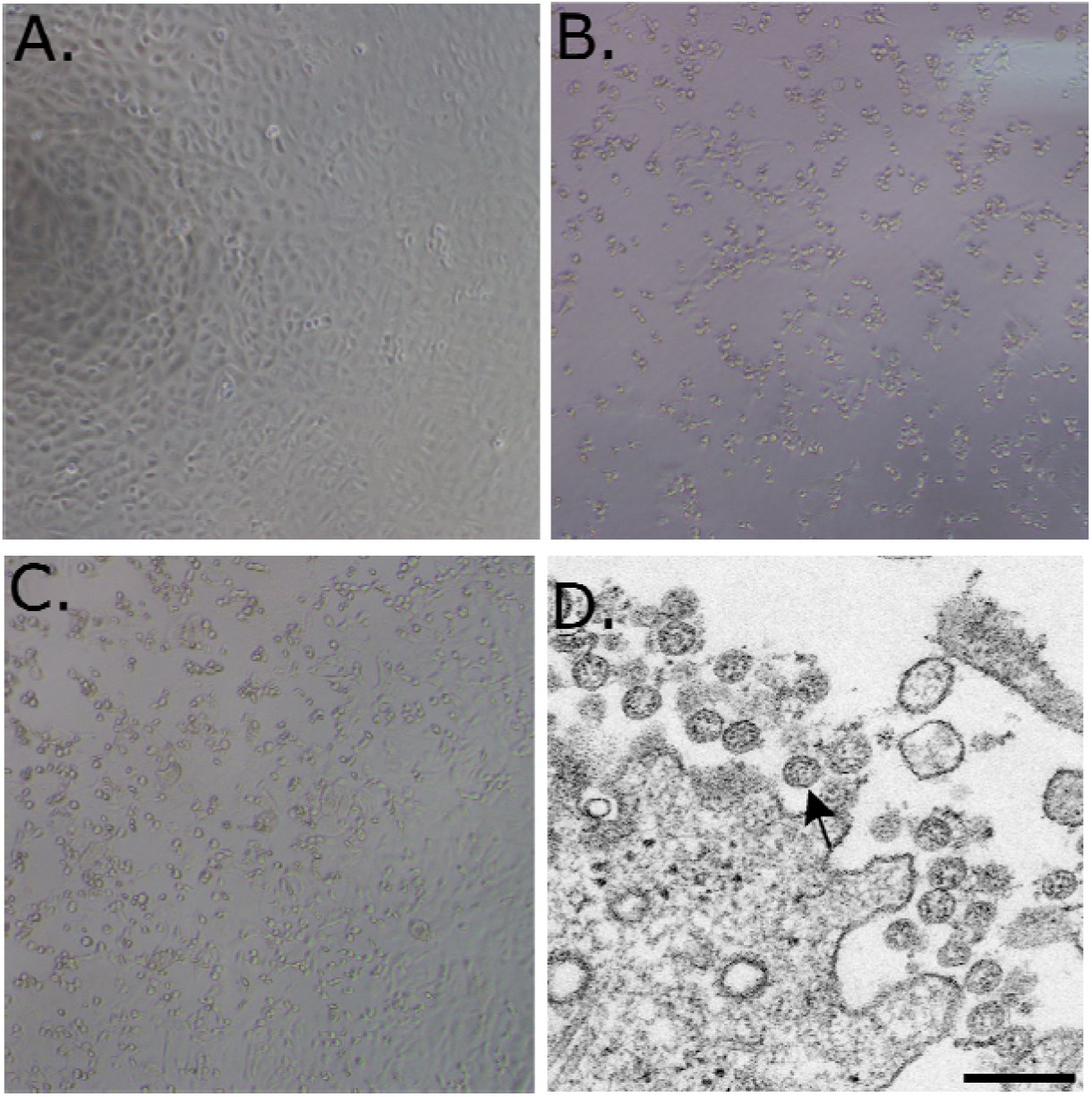
(A-C) 10X phase contrast images of vero monolayers at 3 days post-inoculation. Panels shown are (A) mock, (B) nasopharyngeal, and (C) oropharyngeal. (D) Electron microscopic image of the viral isolate showing extracellular spherical particles with cross sections through the nucleocapsids, seen as black dots.

Isolates from the first passage of an OP and an NP specimen were used for whole genome sequencing. The genomes from the NP specimen (Genbank accession MT020880) and OP specimen (Genbank accession MT020881) matched each other 100%. The isolates also matched the corresponding clinical specimen 100% (Genbank accession MN985325).

After the second passage, OP and NP specimens were not cultured separately. Virus isolate was passaged two more times in Vero CCL-81 cells, and titrated by TCID50. The titers of the third and fourth passages were 8.65 x 10^6^ and 7.65 x 10^6^ TCID50 per mL, respectively.

Of note, we passaged this virus in the absence of trypsin. The spike sequence of SARS-CoV-2 has an RRAR insertion at the S1-S2 interface that may be cleaved by furin (10). Highly pathogenic avian influenza viruses have highly basic furin cleavage sites at the hemagglutinin protein HA1-HA2 interface that permit intracellular maturation of virions and more efficient viral (11). The RRAR insertion in SARS-CoV-2 may serve a similar function.

We subsequently generated a fourth passage stock of SARS-CoV-2 on VeroE6 cells, another fetal rhesus monkey kidney cell line. Viral RNA from SARS-CoV-2 passage four stock was sequenced and confirmed to have no nucleotide mutations compared with the original reference sequence (Genbank accession MN985325). Both SARS-CoV and MERS-CoV had been found to grow well on VeroE6 and Vero CCL81 respectively (12, 13). To establish a plaque assay and determine the preferred Vero cell type for quantification, we titered our passage four stock on VeroE6 and VeroCCL81. Following infection with a dilution series, we found that SARS-CoV-2 replicated in both Vero cell types; however, the viral titers were slightly higher in VeroE6 cells than Vero CCL81 (**Figure 2A**). In addition, plaques were more distinct and visible on Vero E6 (**Figure 2B**). As early as 2 days post inoculations, VeroE6 cells produced distinct plaques visible with neutral red staining. In contrast, Vero CCL81 produced less clear plaques and was most easily quantitated with neutral red 3 days post inoculation. On the individual plaque monolayers, SARS-CoV-2 infection of Vero E6 cells produced cytopathic effect with areas of cell clearance (**Figure 2C**). In contrast, Vero CCL81 had areas of dead cells that had fused to form plaques, but the cells did not clear. Together, the results suggest that VeroE6 may be the best choice for amplification and quantification, but both Vero cell types support amplification and replication of SARS-CoV-2.

**Figure 2.**
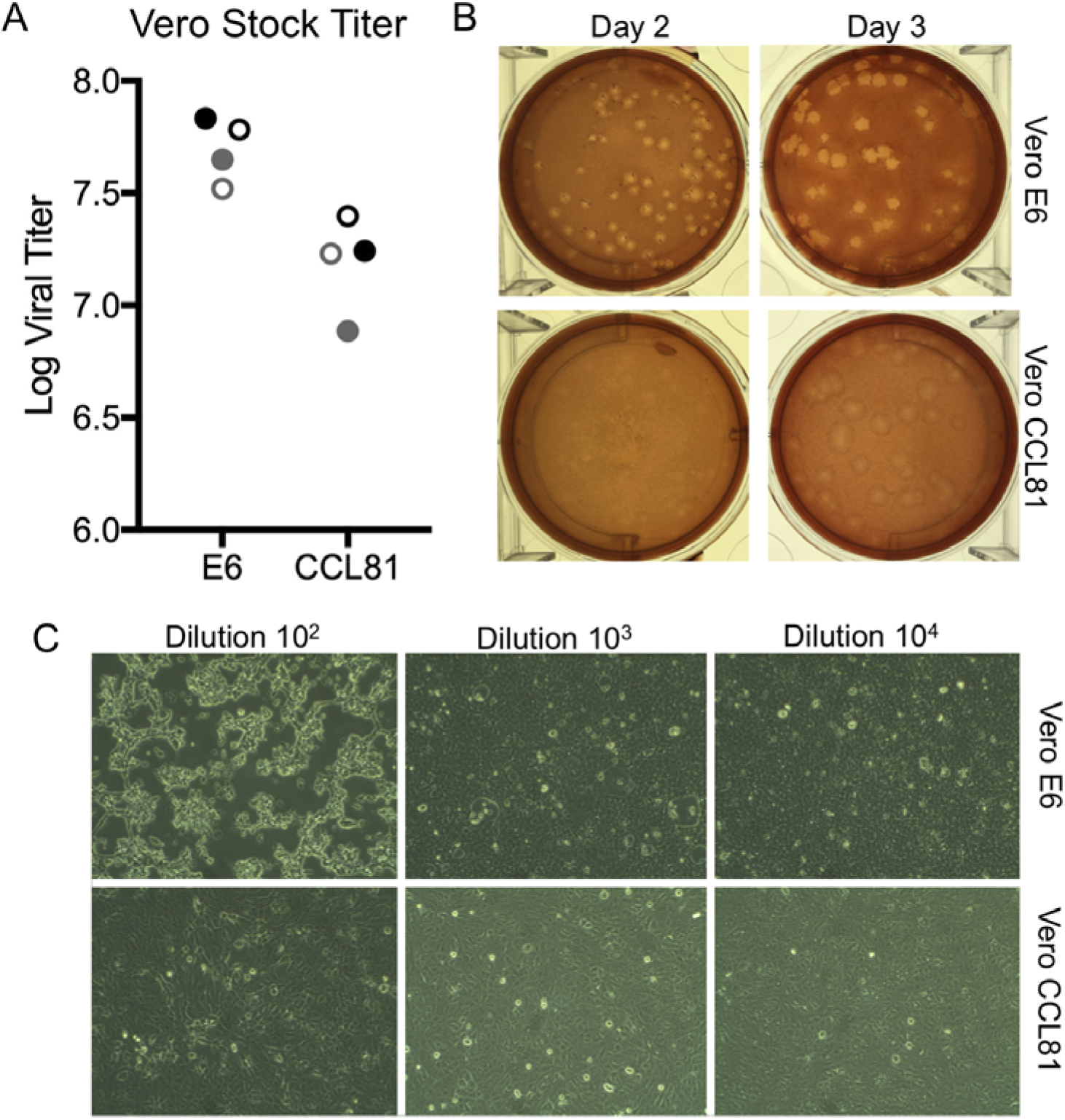
Viral propagation and quantitation. A) Two SARS-CoV-2 passage 4 stocks (black and gray) were quantified utilizing plaque assay at day two (closed circles) and day three (open circles) post infection of Vero E6 and Vero CCL81. B) Plaque morphology on Vero E6 and Vero CCL81 at day 2 and day 3 post inoculation. C) Cell monolayers two days post infection in Vero E6 (top) and Vero CCL81 (bottom row) at multiple dilutions.

As research is initiated to study and respond to SARS-CoV-2, information about cell lines and types susceptible to infection is needed. Therefore, we examined the capacity of SARS-CoV-2 to infect and replicate in several common primate and human cell lines, including human adenocarcinoma cells (A549), human liver cells (HUH7.0), and human embryonic kidney cells (HEK-293T), in addition to Vero E6 and Vero CCL81. We also examined an available big brown bat kidney cell line (EFK3B) for SARS-CoV-2 replication capacity. Each cell line was inoculated with at high MOI and examined 24 hours post infection (**Figure 3A**). No cytopathic effect was observed in any of the cell lines except in Vero cells which grew to >10^7^ PFU at 24 hours post infection. In contrast, both HUH7.0 and 293T cells showed only modest viral replication and A549 cells were incompatible with SARS-CoV-2 infection. These results are consistent with previous susceptibility findings for SARS-CoV and suggest other common culture systems including MDCK, HeLa, HEP-2, MRC-5 cells, and embryonated eggs are unlikely to support SARS-CoV-2 replication (14–16). In addition, SARS-CoV-2 failed to replicate in the bat EFK3B cells which are susceptible to MERS-CoV. Together, the results indicate that SARS-CoV-2 maintain a similar profile to SARS-CoV in terms of susceptible cell lines.

**Figure 3.**
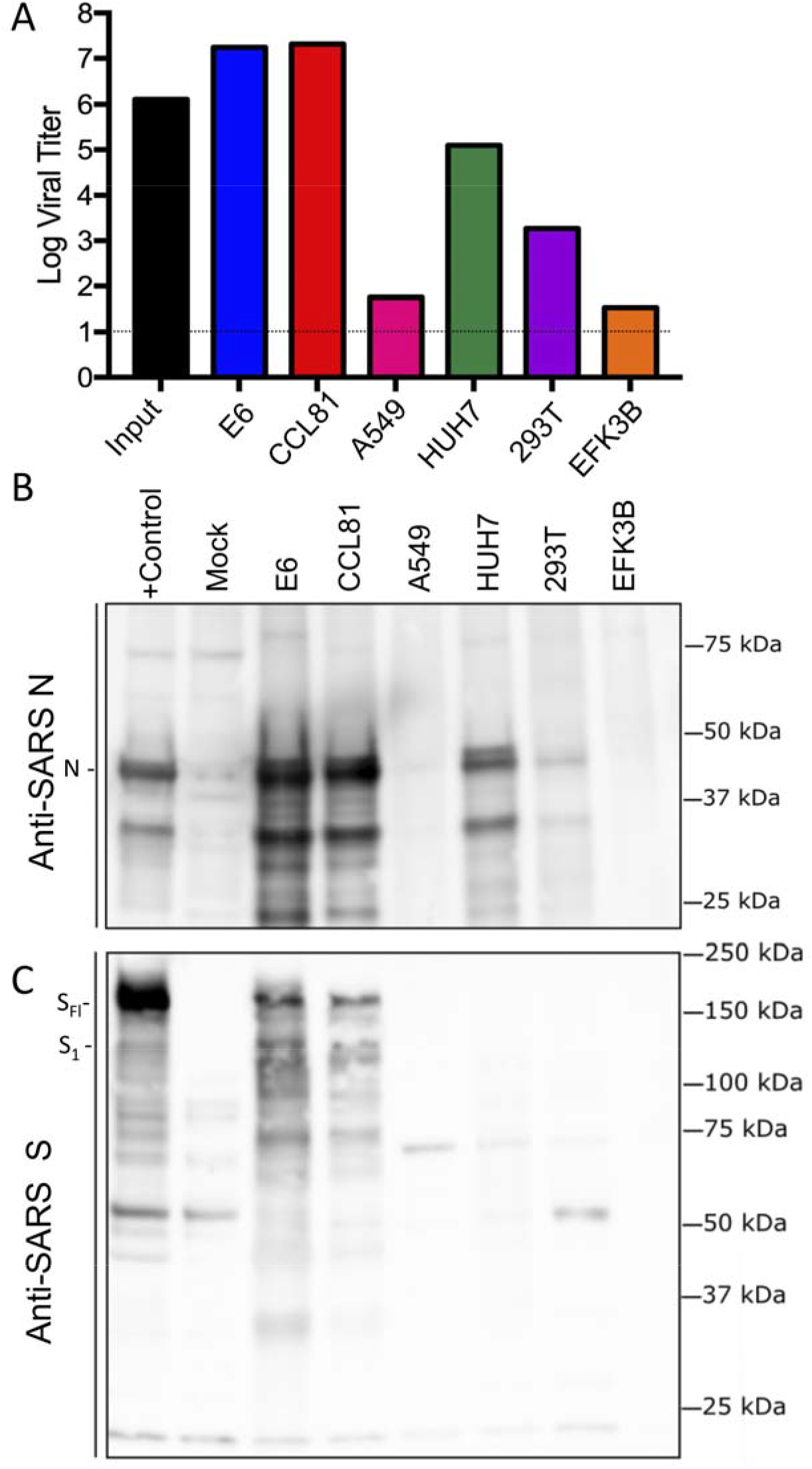
Cell lines susceptible to SARS-CoV-2. (A-C) Cell lines were infected with a high MOI of SARS-CoV-2 (MOI >5), washed after adsorption, and subsequently harvested 24 hours post infection for viral titer and protein lysates. A) Viral titer for SARS-CoV-2 was quantitated by plaque assay on Vero E6 cells two days post inoculation. B & C) Infected cell protein lysates were probed by western blot with B) rabbit polyclonal anti-SARS nucleocapsid (N) antibody or C) anti-SARS-CoV spike (S) antibody. N protein, full length spike (S_FL_) and spike S1 (S_1_) denoted.

Having established robust infection with SARS-CoV-2 in several cell types, we next evaluated the cross reactivity of SARS-CoV antibodies against the SARS-CoV-2. Cell lysates from infected cell lines were probed for protein analysis; we found that polyclonal sera against the SARS-CoV spike and nucleocapsid proteins recognize SARS-CoV-2 (**Figure 3B & C**). The N protein, highly conserved across the group 2B family, retains >90% amino acid identity between SARS-CoV and SARS-CoV-2. Consistent with the replication results (**Figure 3A**), SARS-CoV-2 showed robust N protein in both Vero cell types, less protein in HUH7.0 and 293T, and minimal signal in A549 and EFK3B cells (**Figure 3B**). Similarly, the SARS-CoV spike antibody also recognized SARS-CoV-2 spike protein, indicating cross reactivity (**Figure 3C**). Consistent with SARS CoV, several cleaved and uncleaved forms of the SARS-CoV-2 spike protein. Notably, the cleavage pattern to the the SARS spike positive control from Calu3 cells, a respiratory cell line, varies slightly and could signal differences between proteolytic cleavage of the spike proteins between the two viruses due to predicted insertion of a furin cleavage site in SARS-CoV-2 (10). However, differences in cell type and conditions complicate this interpretation and indicate the need to further study in equivalent systems. Overall, the protein expression data from SARS-CoV N and S antibodies recapitulate replication findings and indicate that SARS-CoV reagents can be utilized to characterize SARS-CoV-2 infection.

Finally, we evaluated the replication kinetics of SARS-CoV-2 in a multi-step growth curve. Briefly, Vero CCL-81 and HUH7.0 cells were infected with SARS-CoV-2 at a low multiplicity of infection (MOI 0.1) and viral replication evaluated every 6 hours for 72 hours post inoculation, with separate harvests in the cell-associated and supernatant compartments (**Fig. 4**). Similar to SARS-CoV, SARS-CoV-2 replicated rapidly in Vero cells after an initial eclipse phase, achieving 10^5^ TCID50 / ml by 24 hours post infection and peaking at > 10^6^ TCID50 / ml. Similar titers were observed in cell-associated and supernatant compartments indicating efficient egress. Despite peak viral titers by 48 hours post-inoculation, significant CPE was not observed until 60 hours post inoculation and peaked at 72 hours post-inoculation, indicating scientists should harvest infected monolayers before peak CPE is observed. Replication in HUH7.0 cells also increased quickly after an initial eclipse phase, but plateaued by 24 hours post inoculation in the intracellular compartment at 2 x 10^3^ TCID50 / ml and dropped off after 66 hours post-inoculation. Virus was not detected in the supernatant of infected HUH7 cells until 36 hours post inoculation and exhibited lower titers at all timepoints (Figure 4). Significant CPE was never observed in HUH7.0 cells. These results are consistent with previous report for both SARS-CoV and MERS-CoV suggesting similar replication dynamics between the zoonotic CoV strains (17, 18).

**Figure 4.**
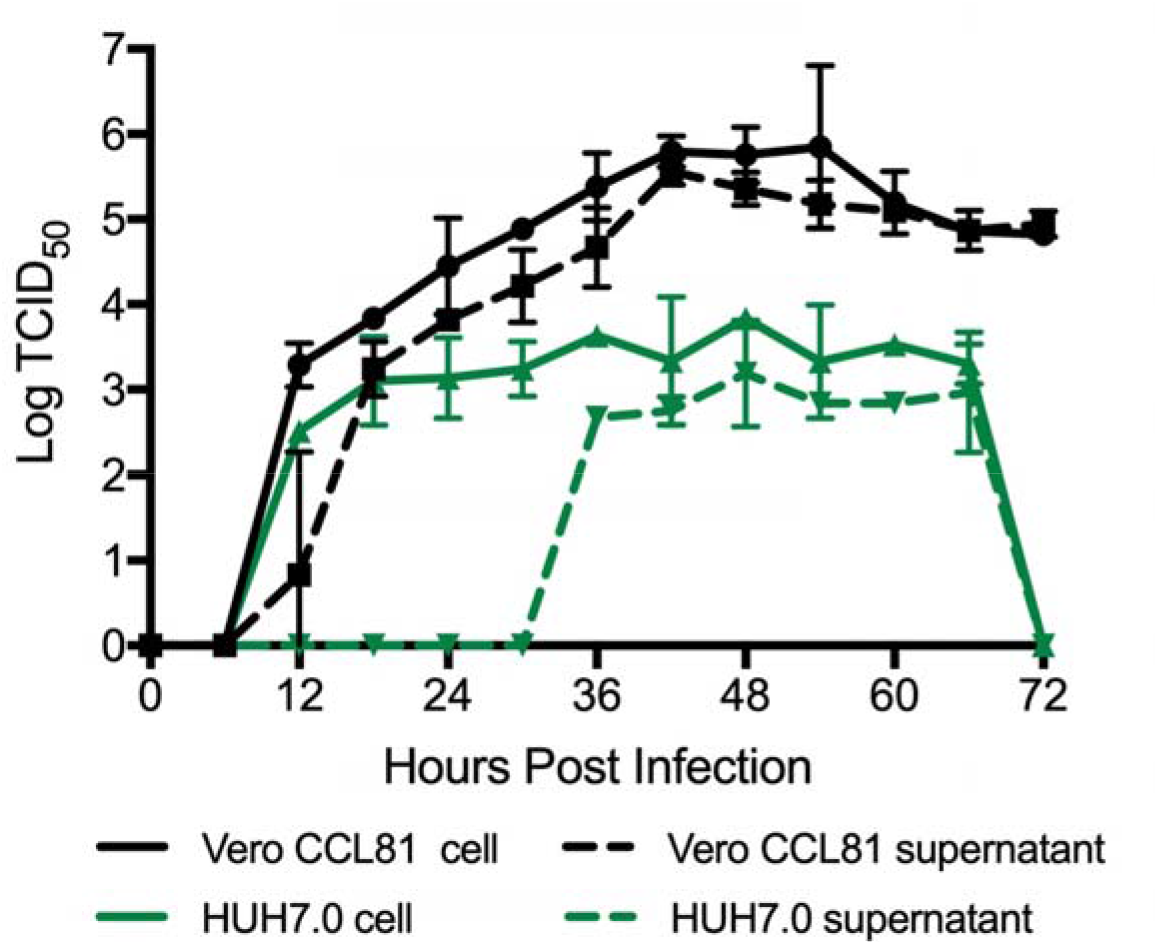
Multi-step growth curve for SARS-CoV-2. Vero CCL81 (Black) and HUH7.0 cells (green) were infected at MOI 0.1 and cells (solid line) and supernatants (dashed line) were harvested and assayed for viral replication via TCID_50_.

The SARS-CoV-2 USA-WA1/2020 viral strain described above has been deposited into BEI reagent resources (ATCC) and the World Reference Center for Emerging Viruses and Arboviruses (WRCEVA, UTMB) to serve as the SARS-CoV-2 reference strain for the United States. The SARS-CoV-2 fourth passage virus has been sequenced and maintains a nucleotide sequence identical to that of the original US clinical strain. These deposits make it available to the domestic and international public health, academic, and pharmaceutical sectors for basic research, diagnostic development, antiviral testing, and vaccine development. We hope broad access will expedite countermeasure development and testing, in addition to facilitating a better understanding of the transmissibility and pathogenesis of this novel emerging virus.

## METHODS

### Specimen collection

Virus isolation from patient samples was deemed to be non-human subjects research by CDC National Center for Immunizations and Respiratory Diseases (research determination 0900f3eb81ab4b6e) Clinical specimens from the first identified US case of COVID-19 acquired during travel to china, were collected as described (1). Nasopharyngeal (NP) and oropharyngeal (OP) swabs in 2 to 3 mL viral transport media were collected on day 3 post-symptom onset for molecular diagnosis and frozen. Confirmed PCR-positive specimens were aliquoted and refrozen until virus isolation was initiated.

### Cell culture, limiting dilution, and isolation

Vero CCL-81 cells were used for isolation and initial passage. Vero E6, Vero CCL-81, HUH 7.0, 293T, A549, and EFKB3 cells were cultured in Dulbecco’s minimal essential medium (DMEM) supplemented with heat inactivated fetal bovine serum(5 or 10%) and antibiotic/antimyotic (GIBCO). Both NP an OP swabs were used for virus isolation. For the isolation, limiting dilution, and passage 1 of the virus, 50 μl serum free DMEM was pipetted into columns 2-12 of a 96-well tissue culture plate. One-hundred μl clinical specimens were pipetted into column 1, and then serially diluted 2-fold across the plate. Vero cells were trypsinized and resuspended in DMEM + 10% FBS + 2X Penicillin-Streptomycin + 2X antibiotic – antimycotic + 2 X amphotericin B at 2.5 x 10^5^ cells / ml. One hundred μl of cell suspension were added directly to the clinical specimen dilutions and mixed gently by pipetting. The inoculated cultures were grown in a humidified 37°C incubator with 5% CO2 and observed for cytopathic effect (CPE) daily. Standard plaque assays were used for SARS-CoV-2 based on both SARS-CoV and MERS-CoV protocols (19, 20).

When CPE were observed, the cell monolayers were scrapped with the back of a pipette tip. Fifty μl of the viral lysate were used for total nucleic acid extraction for confirmatory testing and sequencing. Fifty μl of virus lysate was used to inoculate a well of a 90% confluent 24-well plate.

### Inclusivity / Exclusivity testing

From the wells in which CPE were observed, confirmatory testing was performed using using CDC’s rRT-PCR assay and full genome sequencing (1) The CDC molecular diagnostic assay targets three portions of the N gene, and all three must be positive to be considered positive (https://www.cdc.gov/coronavirus/2019-ncov/lab/rt-pcr-detection-instructions.html) and (https://www.cdc.gov/coronavirus/2019-ncov/lab/rt-pcr-panel-primer-probes.html). To confirm that no other respiratory viruses were present, Fast Track respiratory pathogen 33 testing was performed (FTD diagnostics).

#### Whole genome sequencing

Thirty-seven pairs of nested PCR assays spanning the genome were designed based on the reference sequence, Genbank Accession No. NC045512. Nucleic acid was extracted from isolates and amplified by the 37 individual nested PCR assays. Positive PCR amplicons were used individually for subsequent Sanger sequencing and also pooled for library preparation using a ligation sequencing kit (Oxford Nanopore Technologies), subsequently for Oxford Nanopore MinION sequencing. Consensus Nanopore sequences were generated using minimap 2.17 and samtools 1.9. Consensus sequences by Sanger sequences were generated from both directions using Sequencher 5.4.6, and were further confirmed by consensus sequences generated from nanopore sequencing.

To sequence passage four stock, libraries for sequencing were prepared with the NEB Next Ultra II RNA Prep Kit (New England BioLabs, Inc.) following the manufacturer’s protocol. Briefly, ~70-100 ng of RNA was fragmented for 15 minutes, followed by cDNA synthesis, end repair and adapter ligation. After 6 rounds of PCR the libraries were analyzed on an Agilent Bioanalyzer and quantified by qPCR. Samples were pooled and sequenced with a paired-end 75 base protocol on an Illumina (Illumina, Inc) MiniSeq instrument using the High-Output kit. Reads were processed with Trimmomatic v0.36 (21) to remove low quality base calls and any adapter sequences. The *de novo* assembly program ABySS (22) was used to assemble the reads into contigs, using several different sets of reads, and kmer values from 20 to 40. Contigs greater than 400 bases long were compared against the NCBI nucleotide collection using BLAST. A nearly full length viral contig was obtained in each sample with 100% identity to the 2019-nCoV/USA-WA1/2020 strain (MN985325.1). All the remaining contigs mapped to either host cell ribosomal RNA or mitochondria. The trimmed reads were mapped to the MN985325.1 reference sequence with BWA v0.7.17 (23) and visualized with the Integrated Genomics Viewer (24) to confirm the identity to the USA-WA1/2020 strain.

### Electron microscopy

Infected Vero cells were scraped from the flask, pelleted by low speed centrifugation, rinsed with 0.1M phosphate buffer, pelleted again and fixed for 2 hours in 2.5% buffered glutaraldehyde. Specimens were post fixed with 1% osmium tetroxide, *en bloc* stained with 4% uranyl acetate, dehydrated and embedded in epoxy resin. Ultrathin sections were cut, stained with 4% uranyl acetate and lead citrate, and examined with a Thermo Fisher/FEI Tecnai Spirit electron microscope.

#### Protein Analysis and Western Blot

Cell lysates were harvested with Laemmli SDS-PAGE sample buffer (BioRAD) containing a final concentration of 2% SDS and 5%ß-mercaptoethanol. Cell lysates were the boiled and removed from the BSL3. The lysates were then loaded onto a poly-acrylamide gel, and SDS-PAGE followed by transfer to polyvinylidene difluoride PVDF membrane. The membrane was then blocked in 5% nonfat dry milk dissolved in Tris-buffered saline with 0.1% Tween-20 (TBS-T) for 1 hour, followed by a short TBS-T wash. Overnight incubation with primary antibody, either rabbit polyclonal sera against the SARS-CoV spike (Sino Biological #40150-T52), β-Actin antibody (Cell Signaling Technology #4970), or a custom rabbit polyclonal sera against SARS-CoV nucleocapsid, was then performed. After primary antibody incubation, the membrane was washed 3x with TBS-T, and then horseradish peroxidase-conjugated secondary antibody was applied for 1 hour. Subsequently, the membrane was washed 3x with TBS-T, and incubated with Clarity Western ECL Substrate (Bio-Rad #1705060S), and imaged with a multi-purpose imaging system.

#### Generation of SARS Nucleocapsid antibodies

The plasmid, pBM302 (25), was used to express SARS-CoV N protein, with a C-terminal His6 tag, to high levels within the inclusion bodies of E.coli and the recombinant protein was purified from the inclusion bodies by nickel-affinity column chromatography under denaturing conditions. The recombinant SARS-CoV N protein was refolded by stepwise dialysis against Tris/phosphate buffer with decreasing concentrations of urea to renature the protein. The renatured, full-length, SARS-CoV N protein was used to immunize rabbits to generate an affinity-purified rabbit anti-SARS-CoV N polyclonal antibody.

## ACKNOWLEDGEMENTS

The reagent described is available through BEI Resources, NIAID, NIH: SARS-related coronavirus 2, Isolate USA-WA1/2020, NR-52281. We thank Dr. Mavanur R. Suresh for providing the plasmid, pBM302, expressing the SARS-CoV N protein. Research was supported by grants from NIA and NIAID of the NIH (U19AI100625 and R00AG049092 to VDM; R24AI120942 to SCW; AI99107 and AI114657 to SM). Research was also supported by STARs Award provided by the University of Texas System to VDM, funds from the Institute for Human Infections and Immunity at UTMB to SM, and trainee funding provided by the McLaughlin Fellowship Fund at UTMB

## DISCLAIMERS

The findings and conclusions in this report are those of the author(s) and do not necessarily represent the official position of the Centers for Disease Control and Prevention. Names of specific vendors, manufacturers, or products are included for public health and informational purposes; inclusion does not imply endorsement of the vendors, manufacturers, or products by the Centers for Disease Control and Prevention or the US Department of Health and Human Services.

